## Supplementary Data for "Structure of the human heparan sulfate polymerase complex EXT1-EXT2"

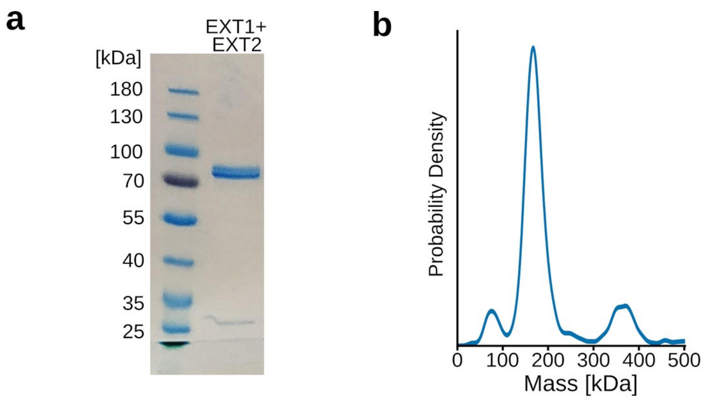


**Extended data Fig. 1 | Purification and functional characterization of the EXT1-EXT2 complex. a**, Coomassie-stained SDS-PAGE analysis of purified EXT1-EXT2 complex, lacking the N-terminal membrane anchoring helix. Expected molecular sizes of EXT1 and EXT2 are about 85 kDa and 79 kDa, respectively. **b**, Mass photometry analysis of purified complex with a major peak at 160 kDa.

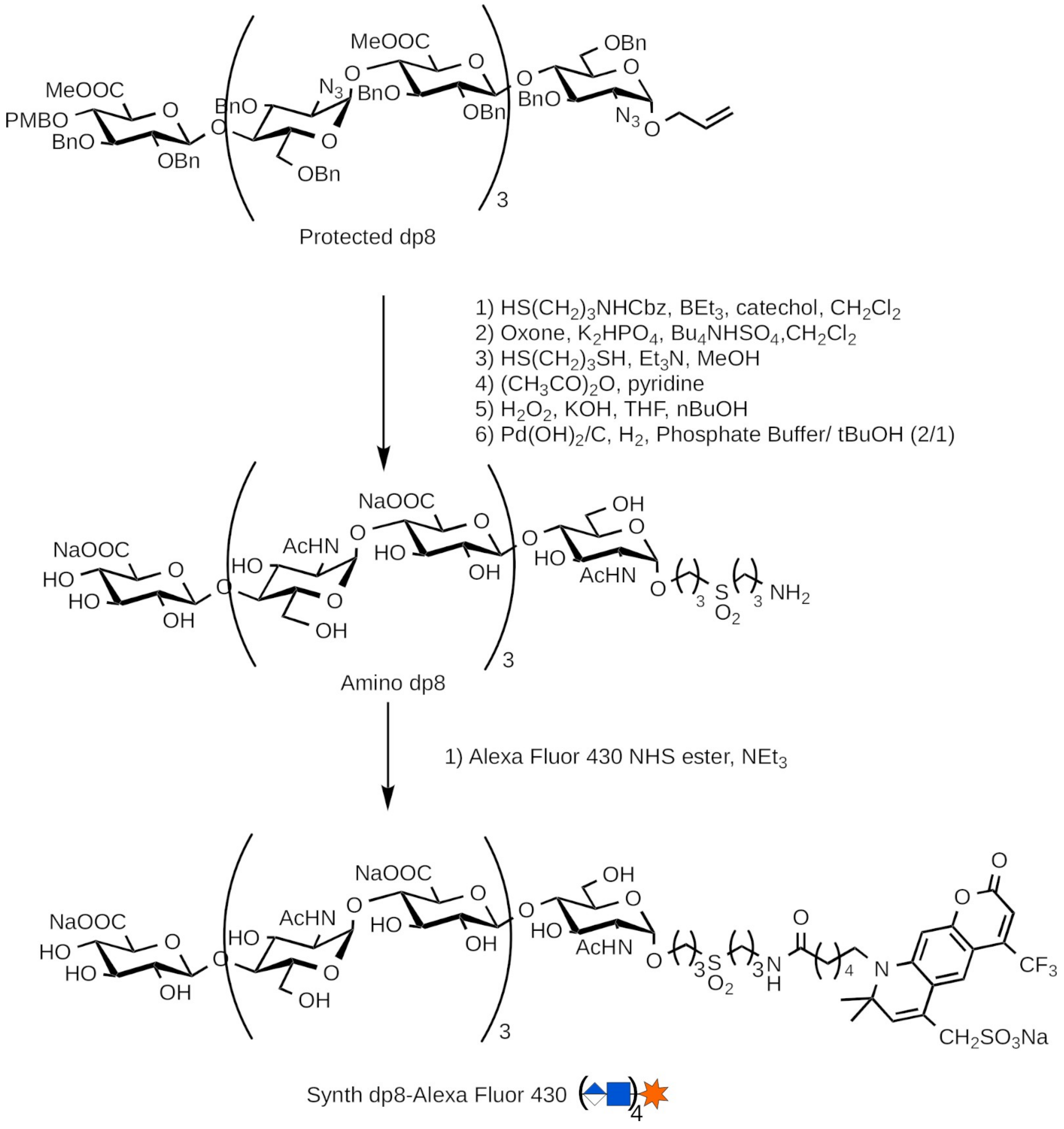


**Extended data Fig. 2: Preparation of the octa-saccharide substrate analog.** 7 step chemical synthesis of synth dp8-Alexa Fluor 430, an octa-saccharide consisting of repeating GlcNAc and GlcA units with a Alexa-fluorophore at the reducing end, from a protected dp8 precursor.

a

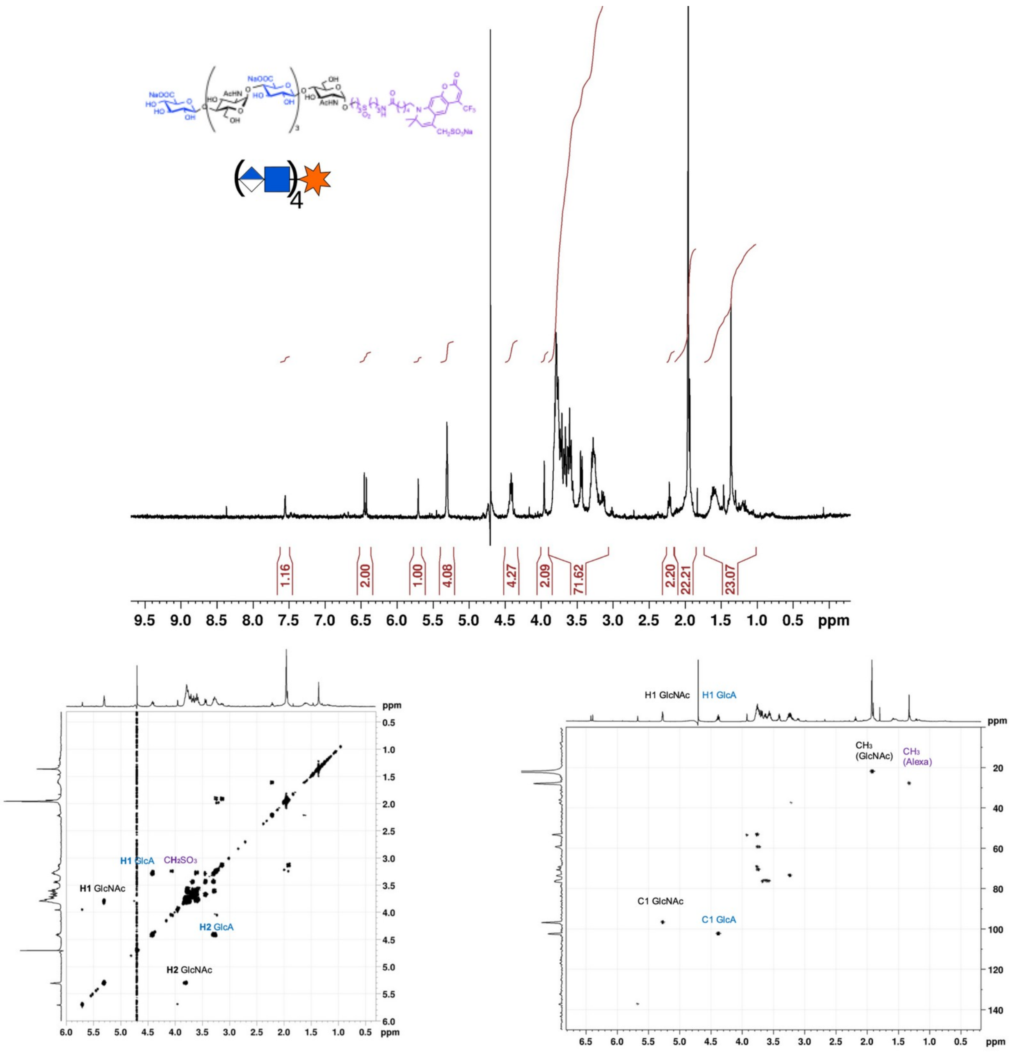


b

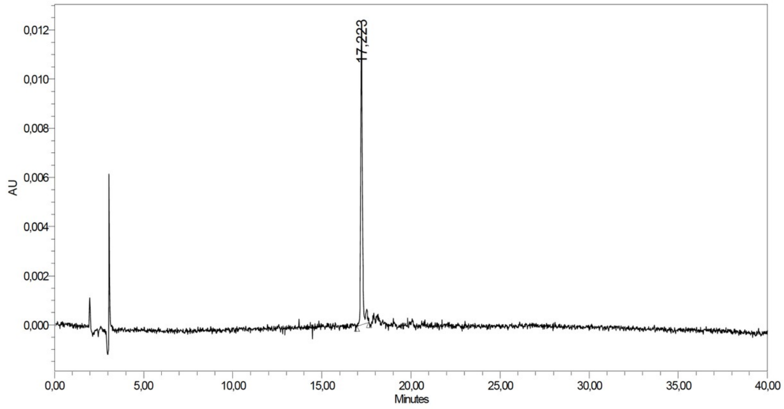


**Extended data Fig. 3: Characterization of the octa-saccharide substrate analog. a**, NMR (1H, COSY, HSQC) spectroscopy of the synth dp8-Alexa Fluor 430. NMR analysis reveals expected signals at characteristic $^{1}H$ and $^{13}C$ chemical shifts of GlcNAc: H1/C1 (5.27/97.4 ppm), H2/C2 (3.77/69.2 ppm), $CH_3$ (1.93/21.8 ppm); GlcA: H1/C1 (4.39/103.1 ppm), H2/C2 (3.25/73.7 ppm); and Alexa Fluor 430: $CH_2SO_3Na$ (3.92/53.8 ppm), $CH_3$ (1.32/28.6 ppm). $^{13}C$ chemical shifts were derived from $^{1}H$-$^{13}C$ HSQC experiment. **b**, HPLC chromatogram of synth dp8-Alexa Fluor 430 indicating 95% purity based on absorbance at 430 nm.

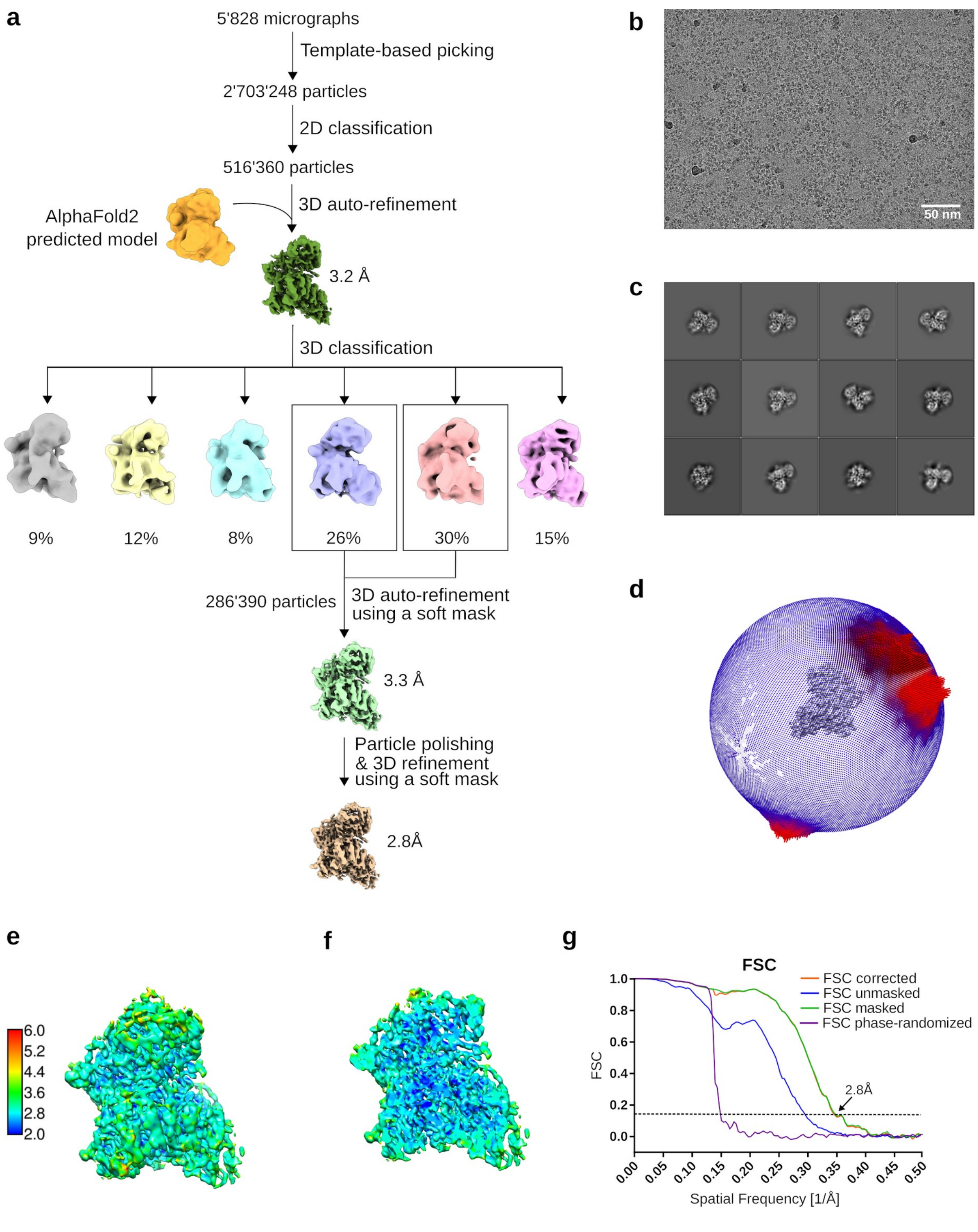


**Extended data Fig. 4: EM data processing and quality assessment for human EXT1-EXT2. a**, Flow-chart of cryo-EM data processing procedure using RELION-3.1. **b**, Representative motion-corrected and dose-weighted micrograph. **c**, Selected class averages from 2D classification. **d**, Angular distribution of particles in the final iteration of 3D refinement. **e**, B-factor sharpened EM map, colored by local resolution as estimated in ResMap. **f**, Map and representation as in (e) but vertically sliced. **g**, Fourier shell correlation (FSC) curve indicating estimated resolutions based on the FSC = 0.143 criterion as generated by RELION-3.1.

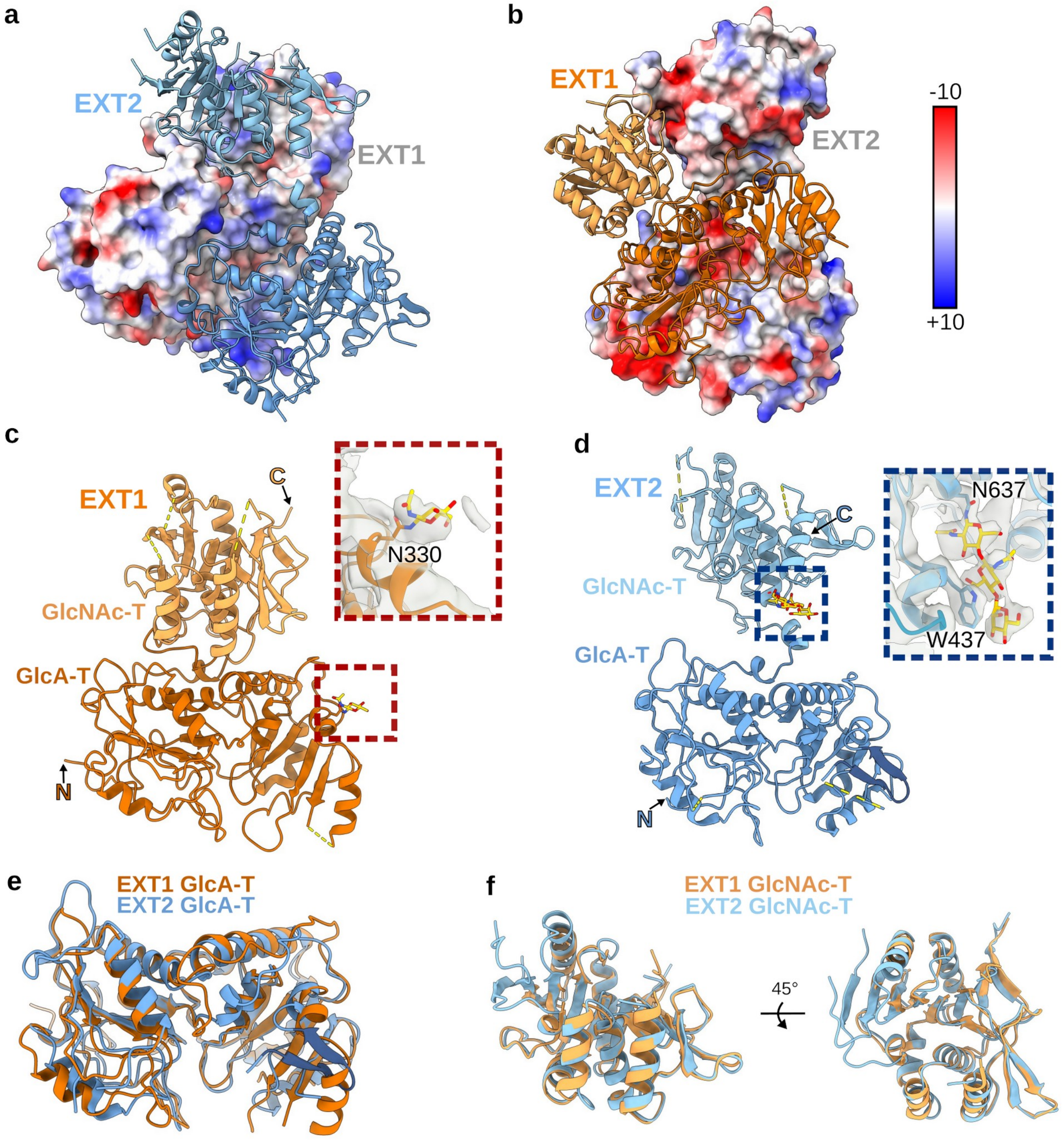


**Extended data Fig. 5 | EXT1 and EXT2 structure comparison. a**, Surface of EXT1 is colored according to its electrostatic potential with isocontours from blue (+10 κT/e) to red (-10 κT/e). EXT2 is shown in ribbon representation with its GlcA-T and GlcNAc-T domain colored in dark and light blue, respectively. **b**, EXT2 is shown in surface representation and colored according to its electrostatic potential. EXT1 is shown in ribbon representation with its GlcA-T and GlcNAc-T domain colored in dark and light orange, respectively. **c**, Cartoon representation of EXT1. Close-up view shows a *N*-glycan in yellow stick representation and surrounding EM map in gray. **d**, Cartoon representation of EXT2. Close-up view shows a well-ordered *N*-glycan, forming a Pi-stacking interaction with Trp437. **e**, Superposition of EXT1 and EXT2 GlcA-T domains with a resulting R.M.S.D of 1.2 Å for 209 of 291 residues. **f**, Superposition of EXT1 and EXT2 GlcNAc-T domains shown from two orientations. R.M.S.D is 0.96 Å for 150 out of 182 aligned residues.

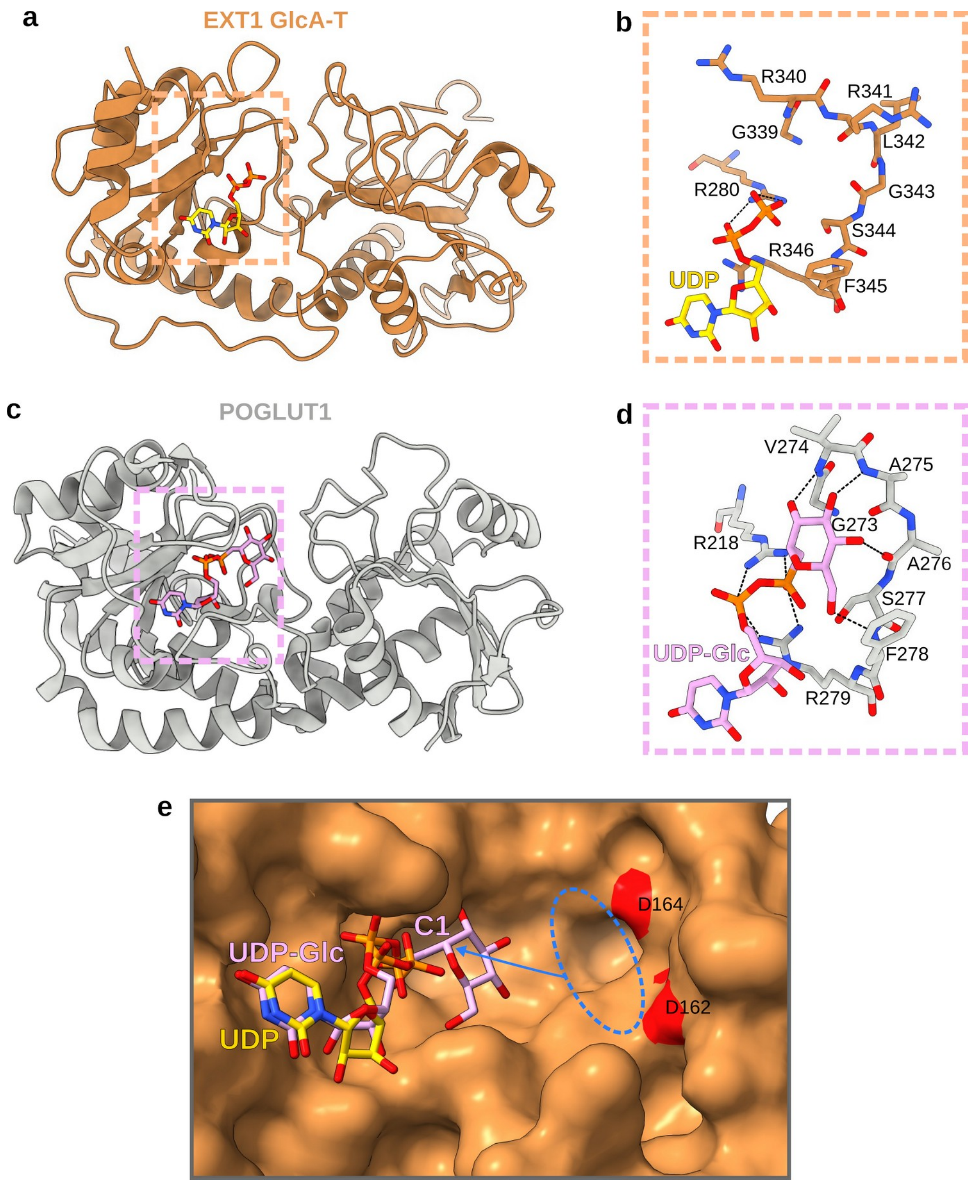


**Extended data Fig. 6 | Substrate binding by EXT1 GlcA-T and human POGLUT1. a**, Cartoon representation of the EXT1 GlcA-T domain shown in dark orange with bound UDP ligand shown as yellow sticks. **b**, Close-up view onto the potential UDP-GlcA substrate recognition loop shown in stick representation. Two critical arginines (R280 and R346) are also shown. **c**, Crystal structure of human POGLUT1 (PDB-ID: 5L0U), a protein *O*-glucosyltransferase, shown in cartoon representation and colored in gray. The bound UDP-Glc substrate analog is shown in stick representation and colored in plumb. **d**, Close-up view shows hydrogen bond network between the protein backbone and the glucose and between arginine residues (R218 and R279) and the two phosphates. The G273-F278 loop accommodating the Glc moiety in POGLUT1 is similar to the loop G339-F345 in the EXT1 GlcA-T domain. **e**, View onto the catalytic site of EXT1 GlcA-T displayed in surface representation with bound UDP ligand shown as yellow sticks. Two aspartate residues potentially involved in acceptor substrate recognition are colored in red. Human POGLUT1 and EXT1 GlcA-T were superimposed and UDP-Glc ligand of POGLUT1 is shown as plum-colored sticks. The potential binding site of the oligosaccharide acceptor substrate is indicated with a blue ellipse and an arrow shows the direction of attack on the C1-carbon, characteristic for a $S_N2$-like reaction mechanism. GlcA, glucuronic acid; Glc, glucose.

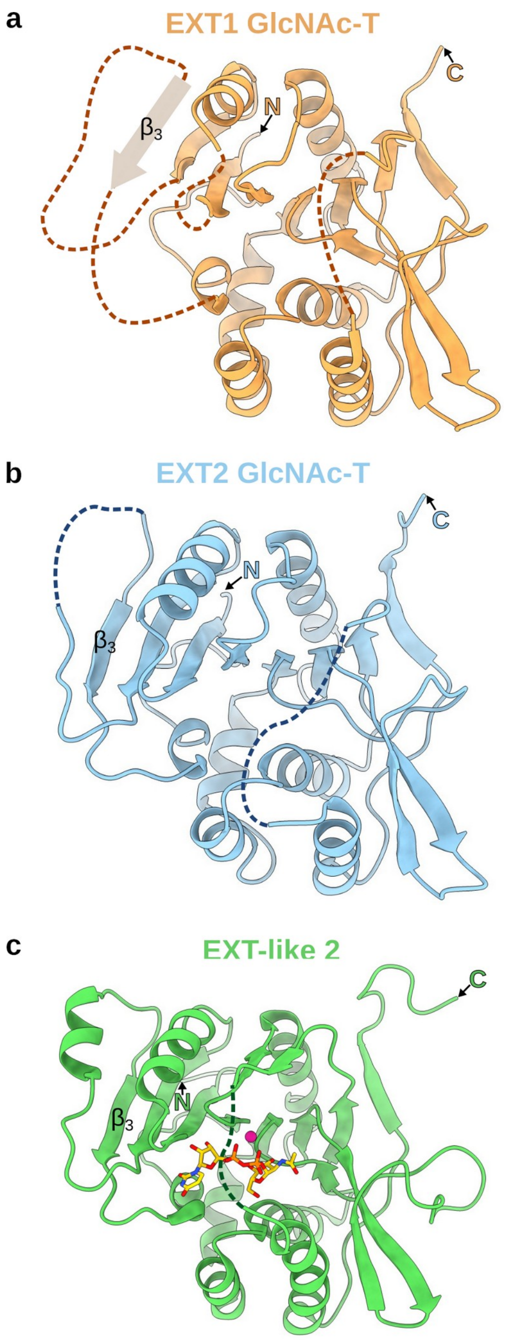


**Extended data Fig. 7 | The architecture of the EXT1 and EXT2 GlcNAc-T domains is highly similar to EXT-like 2. a**, Cartoon representation of the EXT1 GlcNAc-T domain colored in orange. The missing beta-strand $\beta_3$ was drawn by hand. **b**, EXT2 GlcNAc-T domain is shown as cartoon representation and colored blue. **c**, Cartoon representation of the crystal structure of EXT-like 2 (PDB-ID: 1ON6) colored in green with bound UDP-GlcNAc substrate shown as yellow sticks and manganese as pink sphere. Missing loops are indicated as dotted lines.

a

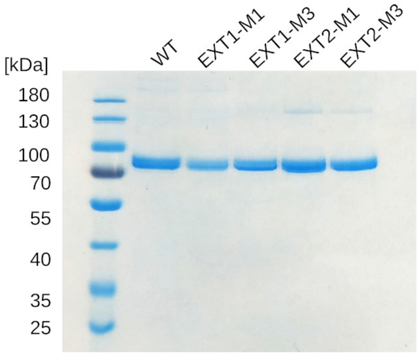


b

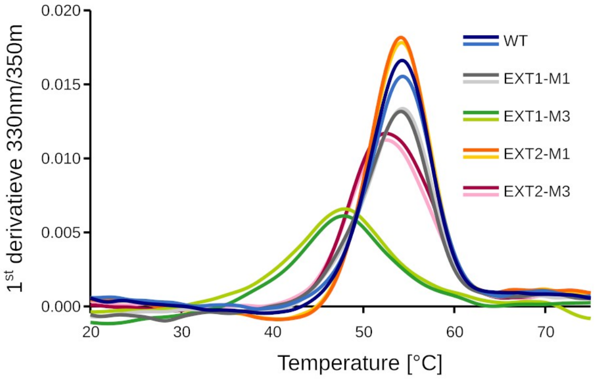


c

| Sample | TM1 [°C] | TM2 [°C] |
|---|---|---|
| WT | 54.2 | 54.3 |
| EXT1-M1 | 54.1 | 54.3 |
| EXT1-M3 | 47.9 | 47.8 |
| EXT2-M1 | 54.1 | 54.2 |
| EXT2-M3 | 52.6 | 52.6 |

d e

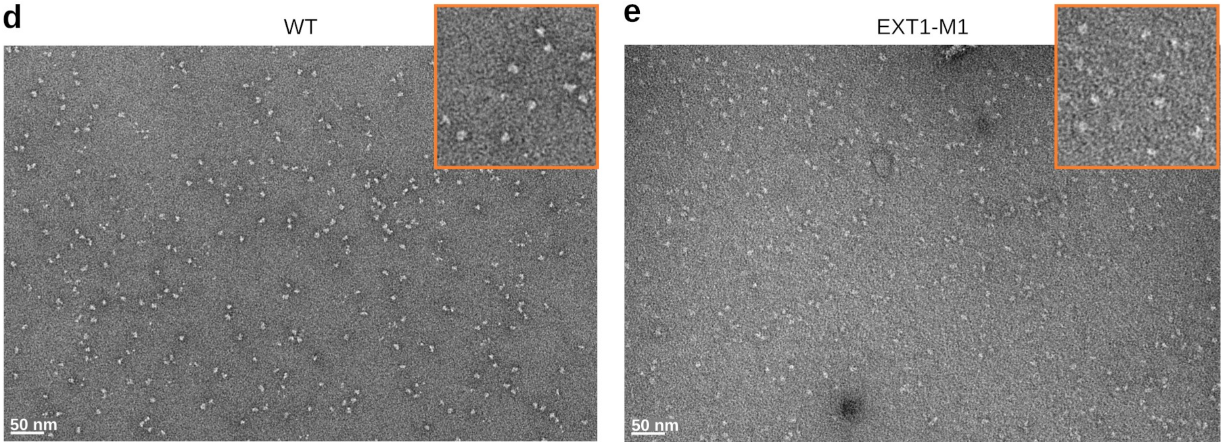


**Extended data Fig. 8 | Quality and stability assessment of purified EXT1-EXT2 complexes harboring point mutations. a,** SDS-PAGE analysis of purified EXT1-EXT2 wild-type (WT) and mutant EXT1-M1 (D162N/D164N), EXT1-M3 (D565N/D567N), EXT2-M1 (D139N/D141N) and EXT2-M3 (D538N/D540N) containing complexes. **b,** Thermal stability measurements of purified EXT1-EXT2 WT and mutant complexes using nano differential scanning fluorimerty. Measurements were performed in duplicate. **c**, Table summarizing the melting temperatures observed in (b). **d-e**, Negative stain EM analysis of the wild-type and EXT1-M1 containing EXT1-EXT2 complex. Zoom-in into an area of the micrograph is shown in the top right corner.

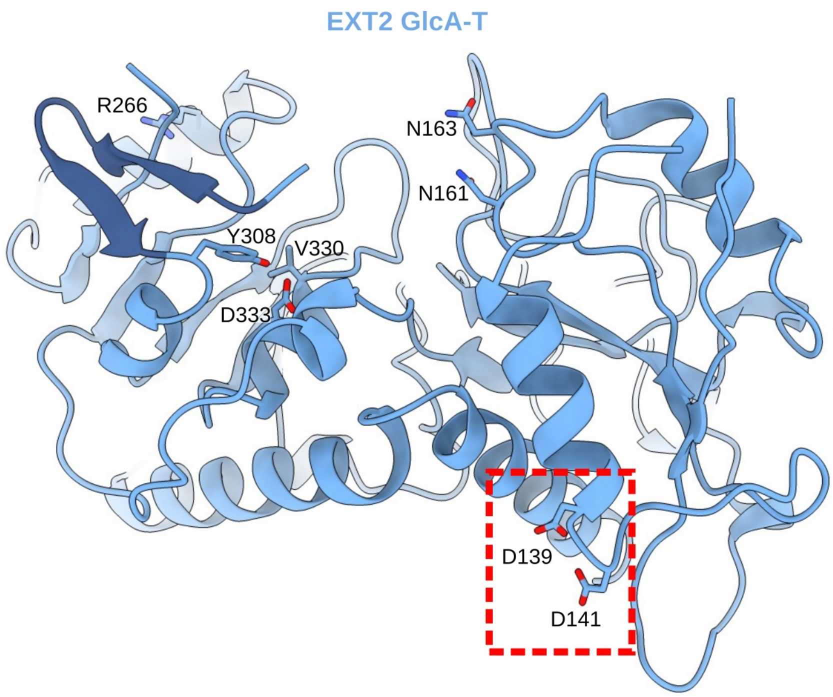


**Extended data Fig. 9 | Location of aspartate residues in EXT2 GlcA-T targeted by mutational analysis.** Cartoon representation of EXT2 GlcA-T with important residues shown in stick representation. Red box highlights a DxD motif, which was replaced during mutational analysis.

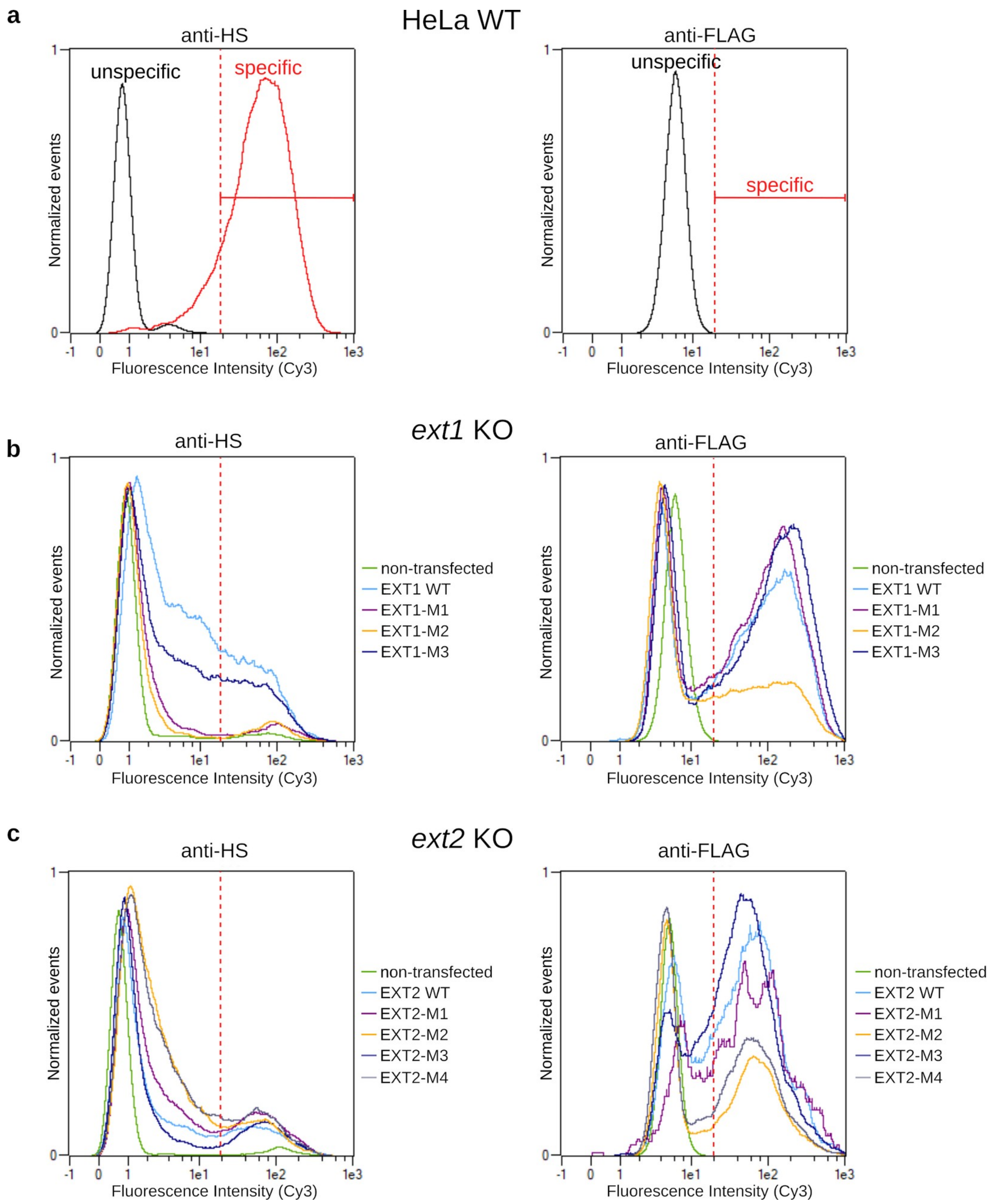


**Extended data Fig. 10 | Flow cytometry analysis of wild-type and knock-out HeLa cells. a**, Flow cytometry plots from a representative experiment, in which 20 000 cells were analyzed. Data was analyzed applying the same threshold (dotted red line), to determine cells giving a HS- or FLAG-tag specific signal, excluding signals such as those from non transfected cells and non-specific binding of antibody. Left panel: wild-type HeLa cells were either labeled with secondary Cy3-conjugated antibody only (unspecific) or with anti-HS and secondary Cy3-conjugated antibody (specific). Right panel: wild-type HeLa cells, not carrying a FLAG-tag, were labeled with anti-FLAG and secondary Cy3-conjugated antibody (unspecific). **b**, Ext1 knock-out HeLa cells (*ext1* KO) were transfected with plasmids encoding wild-type or mutant versions of FLAG-tagged EXT1. Surface HS levels and amount of expressed EXT1 protein were quantified as described in (a). Mutants are described in Figure 3a. **c**, Ext2 KO cell lines were transfected with wild-type or mutant versions of FLAG-tagged EXT2 and HS and EXT2 expression levels were quantified.

**Extended data Table 1: MS-based quantitative characterization of purified human EXT1-EXT2**

| Accession | Protein name | Gene name | MW (Da) | Coverage (%) | Identified peptides | Quantified peptides | iBAQ |
|---|---|---|---|---|---|---|---|
| EXT2 | EXT2 | EXT2 | 78963 | 80.29 | 113 | 109 | 3.04E+08 |
| EXT1 | EXT1 | EXT1 | 85140 | 75.27 | 84 | 83 | 1.77E+08 |
| L0R5A1_HUMAN | Alternative protein CSF2RB | CSF2RB | 11646 | 7.41 | 1 | 1 | 1.54E+07 |
| PPB1_HUMAN | Alkaline phosphatase, placental type | ALPP | 57954 | 64.67 | 35 | 34 | 4.07E+06 |
| B4DGK4_HUMAN | Alternative protein SYT7 | SYT7 | 13173 | 6.56 | 1 | 1 | 2.91E+05 |
| DUS28_HUMAN | Dual specificity phosphatase 28 | DUSP28 | 18324 | 6.25 | 1 | 1 | 9.77E+04 |
| S12A4_HUMAN | Solute carrier family 12 member 4 | SLC12A4 | 120650 | 0.74 | 1 | 1 | 5.01E+04 |
| H2B1A_HUMAN | Histone H2B type 1-A | H2BC1 | 14167 | 7.09 | 1 | 1 | 3.90E+04 |
| SE6L2_HUMAN | Seizure 6-like protein 2 | SEZ6L2 | 97560 | 1.21 | 1 | 1 | 3.87E+04 |
| VIME_HUMAN | Vimentin | VIM | 53652 | 7.73 | 3 | 3 | 3.19E+04 |
| PPBN_HUMAN | Alkaline phosphatase, germ cell type | ALPG | 57377 | 48.68 | 21 | 1 | 1.18E+04 |
| CCNB3_HUMAN | G2/mitotic-specific cyclin-B3 | CCNB3 | 157916 | 0.93 | 1 | 1 | 1.14E+04 |

**Extended data Table 2: EM data collection and structure refinement statistics**

| | |
|---|---|
| **Data collection and processing** | |
| Microscope | Titan Krios |
| Voltage (kV) | 300 |
| Camera | Gatan K3-Summit |
| Energy Filter | Gatan Quantum-LS |
| Magnification | 105’000x |
| Pixel size (Å) | 0.42 |
| Defocus range (µm) | -3.0 to -1.0 |
| Electron exposure ($e^-/Å^2$) | 46 |
| Number of good micrographs | 5’828 |
| Initial number of particles | 2’703’248 |
| Final number of particles | 286’390 |
| Symmetry imposed | No |
| Map resolution (Å) | 2.8 |
| FSC threshold | 0.143 |
| Map resolution range (Å) | 2.5-4.5 |
| **Coordinate and B-factor refinement** | |
| Model resolution (Å) | 2.56 |
| FSC threshold | 0.143 |
| Model resolution range (Å) | 2.56-320 |
| Map sharpening B-factor ($Å^2$) | -30 |
| Number of protein atoms (non-H) | 9’313 |
| Protein residues | 1’146 |
| Number of ligand atoms (non-H) | 78 |
| Mean B-factor protein atoms ($Å^2$) | 71 |
| Mean B-factor of non-protein atoms ($Å^2$) | 78 |
| RMSD bonds (Å) | 0.003 |
| RMSD bond angles (°) | 0.58 |
| Map CC (whole map) – volume? | 0.72 |
| Map CC (around atoms) – mask? | 0.74 |
| **Ramachandran plot** | |
| Favored (%) | 91.4 |
| Allowed (%) | 8.4 |
| Disallowed (%) | 0.2 |
| **Validation** | |
| Molprobity score | 2.14 |
| All-atom clashscore | 13 |
| Rotamer outliers (%) | 0.98 |

**Extended data Table 3: Summary of known miss-sense mutations in EXT1 and EXT2.** Literature was searched for previously described patient mutations which caused hereditary multiple exostoses and which were caused by single amino acid substitutions.

| Protein | Amino Position | Mutation | References |
|---|---|---|---|
| EXT1 | 67 | P→H | 73 |
| EXT1 | 164 | D→H | 74 |
| EXT1 | 220 | S→N | 73 |
| EXT1 | 271 | Y→C | 75 |
| EXT1 | 280 | R→ S | 76 |
| EXT1 | 280 | R→G | 76 |
| EXT1 | 335 | L→R | 77 |
| EXT1 | 339 | G→D | 78 |
| EXT1 | 339 | G→V | 79 |
| EXT1 | 340 | R→H | 73,75,76,79,80 |
| EXT1 | 340 | R→C | 79,81 |
| EXT1 | 340 | R→S | 31 |
| EXT1 | 340 | R→L | 82 |
| EXT1 | 341 | R→G | 83 |
| EXT1 | 346 | R→G | 75 |
| EXT1 | 356 | V→C | 80 |
| EXT1 | 490 | L→R | 80 |
| EXT1 | 644 | N→Y | 73 |
| EXT1 | 739 | G→T | 83 |
| EXT2 | 85 | C→R | 19 |
| EXT2 | 128 | R→W | 73 |
| EXT2 | 152 | L→R | 84 |
| EXT2 | 180 | R→T | 81 |
| EXT2 | 202 | A→V | 85 |
| EXT2 | 223 | R→P | 86 |
| EXT2 | 227 | D→N | 81 |
| EXT2 | 339 | C→Y | 73 |
| EXT2 | 380 | I→T | 87 |
| EXT2 | 576 | E→K | 87 |
